# Improvements to strip-based digital image registration for robust eye-tracking and to minimize distortions in images from scanned ophthalmic imaging systems

**DOI:** 10.1101/2020.12.07.414854

**Authors:** Min Zhang, Elena Gofas-Salas, Bianca T. Leonard, Yuhua Rui, Valerie Snyder, Hope Reecher, Pedro Mecê, Ethan A. Rossi

## Abstract

Retinal image-based eye tracking from scanned ophthalmic imaging systems, such as scanning laser ophthalmoscopy, has allowed for precise real-time eye tracking at sub-micron resolution. To achieve real-time processing rates, strip-based image registration methods for real-time applications have several constraints that limit their performance. This trade-off is acceptable for many imaging and psychophysical applications but when the objective is precise eye motion measurement over time, a high error tolerance can be consequential. Dropped strips in these applications can complicate FEMs quantification. Some light starved imaging applications, such as autofluorescence retinal imaging, also require the retention and registration of as much of the data as possible to increase the signal to noise ratio in the final integrated or averaged image. We show here that eye motion can be extracted from image sequences from scanned imaging systems more consistently when the constraints of real-time processing are lifted, and all data is available at the time of registration. This is enabled with additional image processing steps to achieve a more robust solution. Our iterative approach identifies and discards distorted frames, detects coarse motion to generate a synthetic reference frame and then uses it for fine scale motion tracking with improved sensitivity over a larger area. We demonstrate its application here to tracking scanning laser ophthalmoscopy (TSLO) and adaptive optics scanning light ophthalmoscopy (AOSLO). We show that it can successfully capture most of the eye motion across each image sequence, leaving only between 0.04-3.39% of non-blink frames untracked, even with low quality images, while simultaneously minimizing image distortions induced from eye motion. These improvements will facilitate precise FEMs measurement in TSLO and longitudinal tracking of individual cells in AOSLO.

## I. INTRODUCTION

Fixational eye movements (FEMs) are an essential aspect of normal human vision that keep the eyes in constant motion^1^. FEMs are physiologically important and serve several useful purposes^1–3^. However, they are a source of image distortions in ophthalmic instruments that scan an imaging beam across the retina of awake human observers, such as scanning laser ophthalmoscopy (SLO) and optical coherence tomography (OCT). Scanned systems typically scan the retina in a raster pattern implemented with a relatively slow scanner at the frame rate and a faster scanner at the line rate. SLO line rates are often on the order of ∼10–14kHz, so each line is usually considered to be free from eye-motion based distortions. However, intra-frame image distortions arise because eye movements can be much faster than the relatively slow frame rate of scanned ophthalmic imaging systems that typically operate at video rates (∼24–30Hz). Fortunately, when image sequences are acquired on scanned systems, the motion of the eye is encoded into the image sequence, permitting the motion to be recovered through the application of appropriate techniques^4,5^.

Several methods have been developed to recover eye motion from retinal images acquired with scanned ophthalmic systems, such as SLO^5–7^ and its higher-resolution implementation in adaptive optics scanning laser ophthalmoscopy (AOSLO)^4,8^. Recovery of eye motion for image registration is essential in ophthalmic imaging so that several images from a sequence can be integrated or averaged to generate a high signal to noise ratio (SNR) image of the retina from a sequence of low SNR images. Scanned ophthalmic image registration algorithms have often used a cross-correlation method to compute the offset between a reference image and each image (or sub-image) in an image sequence. To increase the temporal rate of motion measurement, reduce computational cost, and achieve superior results for image registration, most implementations now divide each frame into multiple image strips with a height of several scan lines^9^.

Strip-based image registration methods that track motion based on using a single imaging frame as a reference image have successfully achieved high performance at real-time data rates and enabled previously unattainable engineering and scientific aims, such as real-time optical stabilization ^9,10^ and single-cell psychophysics in AOSLO^11,12^. However, this high performance for real-time applications comes at the cost of a high error tolerance and a reduced sensitivity to large amplitude motion. A high error tolerance results in either dropped strips or poor registration matches. A high error tolerance is acceptable (and sometimes advantageous) for some retinal imaging and most psychophysical applications. Psychophysical applications often tend to consist of hundreds or thousands of trials, so can usually simply just exclude those trials from analysis when the algorithm dropped a strip or gave a bad match. Poor-matches can occur when the image is of poor quality due to optical factors, so the dropping of these poor-quality strips can be advantageous when the objective is to build a registered and averaged image from only the highest quality image strips (e.g. confocal AOSLO). However, the same error-tolerance is unacceptable when the objective is precise eye motion measurement (e.g. to quantify fixational eye movements for tens of seconds or up to minutes), or in cases with low photon flux (e.g. in autofluorescence or non-confocal AOSLO).

A drawback to the single reference frame approach is that it is insensitive to motion that moves the field of view outside the area imaged in the reference frame; this is sometimes referred to as a ‘frame-out’ error. We and our colleagues previously demonstrated methods to increase sensitivity to large amplitude motion for eye tracking in AOSLO using hybrid imaging approaches^9,13^. These add an additional wide-field scanned system to track large amplitude motion in real-time at a coarse scale to drive a tip/tilt mirror to stabilize the small field of view AOSLO enough to prevent most ‘frame-out’ errors. However, these solutions increase the complexity of the imaging system and the data acquisition and image processing pipelines substantially.

Finally, previous approaches that used only a single reference frame do not mitigate or remove the intraframe distortions present within the reference frame but rather encode the reference frame distortion into all the images in the registered image sequence. Some methods have been proposed to mitigate these distortions, but no satisfactory solution has been shown to routinely reconstruct the same spatial arrangement between image features independent of the starting reference frame. Though within-frame distortions introduced from the reference frame are usually small (on the order of several microns), they have historically made it extremely difficult to track individual cells longitudinally in AOSLO^14^.

Here we show an improved approach that solves these problems for strip-based registration and demonstrate its applications in TSLO and AOSLO. Free from the constraints imposed by real-time eye-tracking, we devised a more robust approach that achieves our goals for a technique that: 1) tracks the precise motion of nearly all the images in each sequence for eye-tracking and light starved imaging applications; 2) is sensitive to motion larger than the field of view of a single frame; and 3) reconstructs the spatial arrangement between image features consistently and accurately. Our approach also applies some pre-processing steps to achieve robust eye tracking in cases of large variations in illumination with gaze position, such as can arise in TSLO when imaging younger eyes due to the foveal reflex.

We show here applications of the technique for tracking FEMs with TSLO and for image registration in AOSLO and compare the results of this method to the real-time method of Yang et al^9^. Our approach was able to track around 99% of all image strips, on average, after excluding blinks and even in the cases of subjectively lower quality data for both AOSLO and TSLO. We also demonstrate that image distortions are substantially minimized with our new technique.

## II. METHODS

### Participants

All experiments were approved by the University of Pittsburgh Institutional Review Board and adhered to the tenets of the Declaration of Helsinki. Written informed consent was obtained from all participants following an explanation of experimental procedures and risks both verbally and in writing. Participants ranged in age from 14 to 59 (average: 20; female: 20; male: 21) and were compensated for their participation. To ensure that imaging was safe, all light levels were kept below the limits imposed by the latest ANSI standard for safe use of lasers^15^.

### Data sources

We evaluated our approach using sixty image sequences that had been previously acquired on our TSLO and AOSLO systems for ongoing studies^16,17^. Since we were interested in evaluating performance on a range of image qualities, we selected ten image sequences from each system in three general levels of image quality. Images were graded subjectively by two of the authors (MZ and EG) to be of either low, medium, or high quality; both graders had to agree on the grading for the image sequence to be included in the evaluation dataset. Examples of each quality level for each device are shown in supplementary figure 1.

#### Tracking scanning laser ophthalmoscopy (TSLO)

The tracking scanning laser ophthalmoscope (C. Light Technologies, Inc., Berkeley, CA) has been described in detail^10^. An 840 nm (50 nm bandwidth) super luminescent diode (SLD) provided illumination over a field size of 5°×5°. Participants were imaged sitting in a chin rest that was stabilized with temple pads. Image sequences (512×512 pixels) were acquired monocularly, from the left eye, without dilation, at 30 Hz for 30 seconds.

#### Adaptive optics scanning laser ophthalmoscopy (AOSLO)

The Pittsburgh adaptive optics scanning laser ophthalmoscope has been described in detail elsewhere^17^; image sequences were used from the confocal imaging channel only. A 795 nm (FWHM = 15 nm) SLD provided illumination over a field size of 1.5°×1.5°. Image sequences (512×496 pixels) were acquired monocularly, from either the left or right eye, with dilation, at 29 Hz for 10–180 seconds.

### Algorithm workflow

The algorithm workflow, implemented in Matlab (R2018a; The MathWorks Inc., Natick, MA), is outlined in Figure 1. It consists of three main steps: 1) pre-processing and distortion frame detection; 2) coarse registration, large motion detection and synthetic reference frame generation; and 3) fine strip-level registration. The overall strategy follows the general framework described by Stevenson and Roorda^4^ and that we and our colleagues have built upon^8–10^. However, we have added important additional key steps to achieve the goals outlined previously. Each step is described in detail in the sections below. Our workflow begins with blink detection followed by a pre-processing stage.

**Figure 1.**
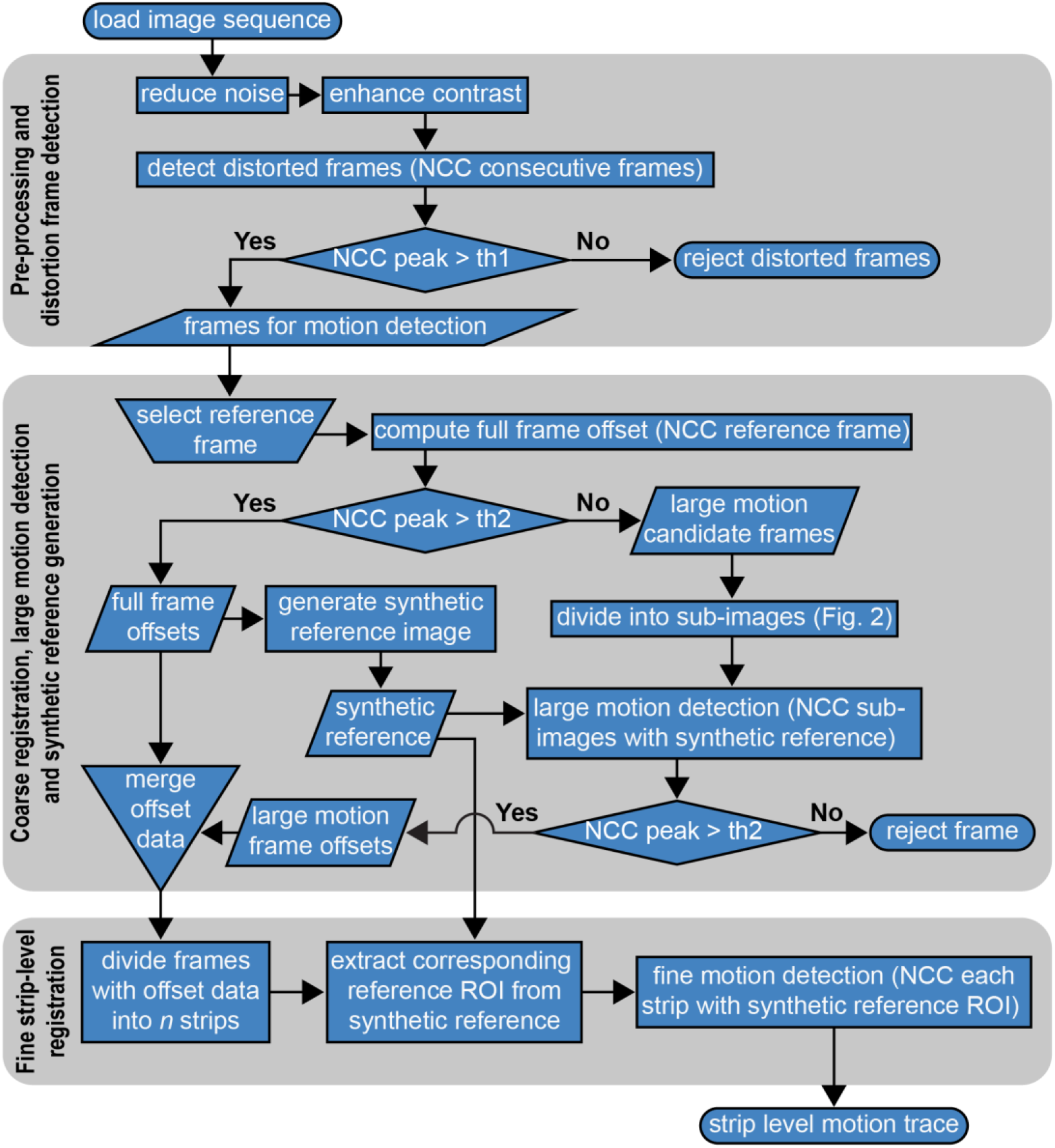
Algorithm workflow. An image sequence is loaded from the database to begin the pre-processing and distortion frame detection stage (top grey rectangle). Noise reduction and contrast enhancement are followed by the detection of distortion frames (NCC on consecutive frames). Frames below threshold 1 (th1) are rejected as distorted frames. Frames above th1 are passed to the next stage for coarse registration, large motion detection and synthetic reference frame generation (middle grey rectangle). A reference frame is selected manually, followed by a full frame offset NCC calculation using the manually selected reference. Frame offsets detected with an NCC peak greater than threshold 2 (th2) are applied to generate the synthetic reference image. Those frames below th2 are divided into sub-images (see figure 2) for large motion detection (NCC between sub-images and synthetic reference). Those sub-images with an NCC peak below th2 are discarded while those above th2 are used to detect the large motion offsets that are merged with the offset data used to create the synthetic reference. The merged offset data is then passed to the final stage of processing, fine strip-level registration (bottom grey rectangle). Strip-level registration is computed by dividing each frame with offset data into *n* strips. The full frame offset data is used to determine the position of the corresponding reference ROI on the synthetic reference. Fine-scale motion is detected by computing the NCC between the reference ROI and the strip (see Fig. 3). A strip-level motion trace is then output.

#### Blink detection

Blinks were detected using a rudimentary intensity threshold to logMean(i), where i ∈[1,2,…,m] is the index of each image and m is the total number of images in the sequence. LogMean(i) is computed by normalizing the mean intensity of each image (i) to the interval of [0, 1], followed by applying the common logarithm on the normalized mean. Note that NaN will be used if the common logarithm of 0 is encountered and hence will not be considered in the threshold calculation. The threshold was defined as: blinkth = minimum(logMean(i))/3.5. Images with logMean ≤ blinkth are marked as blinks and excluded from further processing. It should be noted that this method will erroneously detect blinks for motion traces that do not contain a blink.

#### Pre-processing

Pre-processing reduces noise, improves contrast, and minimizes large gradients in intensity across the field of view. Non-uniform image intensity across the field of view can result from highly scattering structures in the normal retina, such as the foveal reflex; they can often also arise anywhere in the retina in disease states such as age-related macular degeneration. A strong foveal reflex is often seen in younger healthy eyes and was observed in much of our TSLO data. Pre-processing started with Gaussian filtering, implemented with Matlab’s built-in `imgaussfilt` function (with σ = 20), to remove high-frequency noise. This was followed by contrast-limited adaptive histogram equalization (CLAHE) using the `adapthisteq` function (with ClipLimit = 0.05). CLAHE serves to effectively improve local image contrast. Distorted images were identified by computing the normalized cross-correlation (NCC) for each pair of consecutive images in each sequence, after Salmon and colleagues^18^. All NCC computations were performed with Matlab’s built-in `normxcorr2` function, using the CUDA implementation with elements of the Parallel Computing Toolbox. Registration was carried out on machines equipped with Nvidia GPUs (GTX 1080). Since highly distorted images have little or no overlapping features with both the previous and consecutive frame in the sequence, the peak of the NCC matrix computed between these frames is low. We identified candidate distortion frames based on this principle by applying a simple threshold based on the statistics of the image sequence. When the peak in the NCC matrix between the previous frame and consecutive frame were both less than the threshold, we considered these frames to be distortion frames. The threshold was defined as NCC < µ - 0.8δ, where µ and δ are the average and SD of the NCC peak for the entire image sequence. Distortion frames were excluded from further analysis.

#### Coarse alignment and identification of candidate ‘large motion’ frames

This step computes coarse frame-to-frame offsets and synthesizes a composite reference frame for fine motion extraction. This begins with the selection of a reference frame by the experimenter (e.g. Fig 2a). The full-frame NCC is then computed between each of the non-distorted frames and the reference frame, as we and others have described previously^9–11,19^. In practice, some image frames may have a small overlap with the reference frame due to relatively large eye motion (e.g. 2c), resulting in a reduced NCC peak. To maximize the chances that these frames could be captured in our analysis, we set a second threshold at µ - 0.6δ to extract these frames as a group of candidates of ‘large motion’ frames; large motion frame candidates were then reserved for additional processing steps (described below).

**Figure 2.**
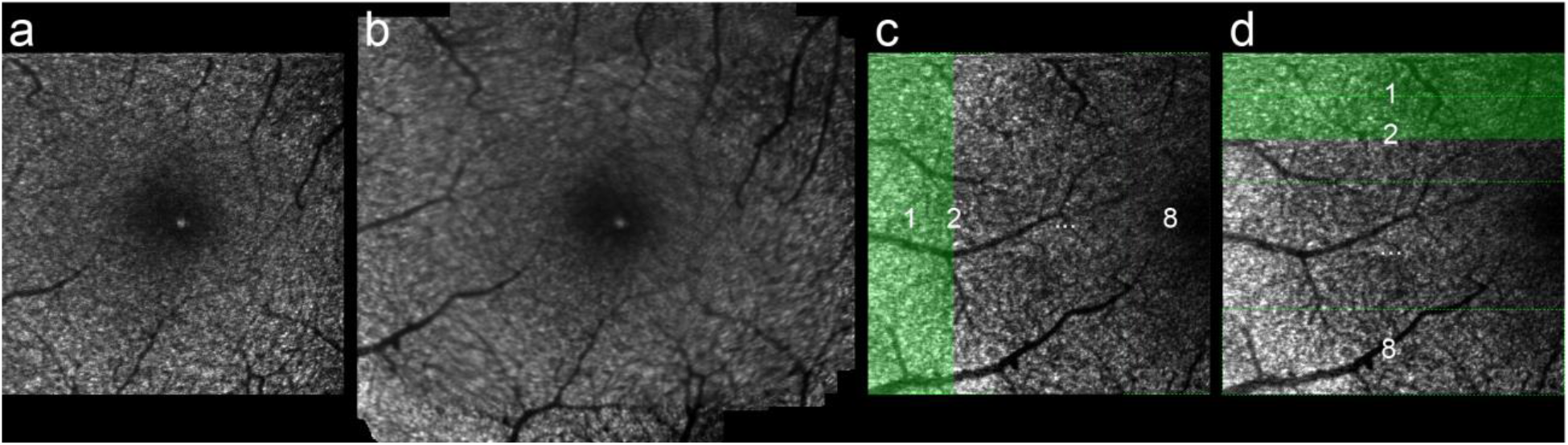
Coarse alignment for reference synthesis and detection of ‘large motion’ candidates. An image from the sequence is chosen manually to serve as the reference for coarse alignment (a). Coarsely aligned frames are averaged to generate a synthetic reference frame (b). Large candidate motion frames (e.g. c and d) are divided into 8 strips both vertically (c) and horizontally (d) and cross-correlated with the synthetic reference frame to capture large amplitude motion.

### Generation of a synthetic reference frame and evaluation of large motion candidates

The next step was to synthesize a larger reference frame for the fine motion trace computation. A larger reference frame enables motion that goes beyond the bounds of a single reference frame to be captured. This was generated by simply averaging the registered images from the coarse alignment step (see Fig. 2b). Next, we attempted to capture the coarse offsets for the images that we determined to be large eye motion candidates. We did this by dividing each of those frames into seven strips vertically and horizontally (see Fig. 2c and 2d). Strip size for this procedure was 512×128 pixels or 128×512 pixels, with 64 pixels of overlap. We then calculated the NCC between each strip and the synthetic reference image. The offset of the strip with the highest NCC peak was selected to represent the true offset of the corresponding frame. If the NCC peak of all the strips for that frame was still low (< µ - 0.6δ), the image was rejected from further analysis.

### Fine motion trace extraction

In the previous step we identified and flagged distortion frames, computed frame-to-frame offsets, generated a composite reference frame and evaluated frames with the largest motion. In this step, we divide each image whose coarse motion was successfully captured, into multiple image strips as shown in Figure 3. The strip size and overlap can be specified by the user; we used 16 strips per image (i.e. 32×512 pixels) herein for a nominal temporal sampling rate of 480 Hz for the motion traces. For each strip, we identified its coarse matching region in the synthetic reference frame from its coarse frame offset. We then extract a larger region of interest (ROI) reference strip by extending the region one strip height above and one strip below the coarse matching region (Fig. 3a). We then use this ROI reference strip for the NCC computation for that strip rather than the whole synthetic reference frame. This approach reduces the computation cost and has the potential to increase accuracy, since the best matching location is usually within the ROI reference strip.

**Figure 3.**
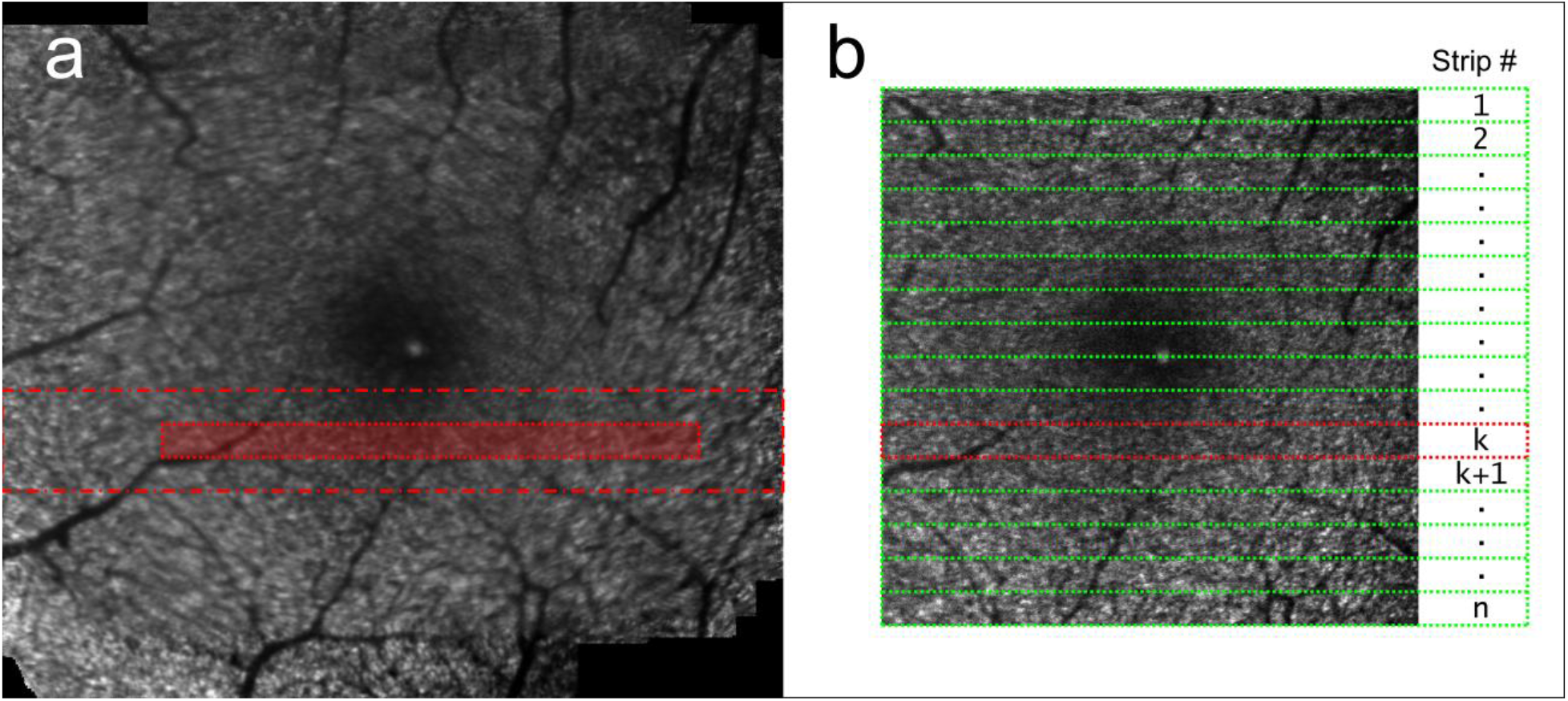
Strip-level motion trace extraction. The synthetic reference image (a) serves as the source of reference ROIs for fine strip-level motion extraction. Each target frame (b) is broken up into strips (e.g. strip k, outlined in red in b) and its probable position on the synthetic reference (shaded area in a) is estimated from its coarse frame offset. A region of interest (ROI) reference area (denoted by the dashed red rectangle in (a)) is then defined and fine motion is extracted by computing the NCC between each strip and its corresponding reference ROI area.

For the example shown in figure 3, if Y_frm_offset_(i) and X_frm_offset_ (i) denote the frame offset of the raw target frame (Fig. 3b), and the region of its kth strip is [x_k_, y_k_, w, h], where (x_k_, y_k_) is the starting position of the strip, w is the width and h is the height of the strip (in our case w = 512 pixels, and h = 32 pixels), then its matching region in the reference frame (shaded red area in fig. 3a) can be determined by [x_ref_, y_ref_, w, h], where

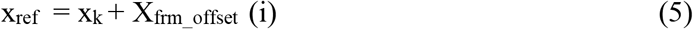

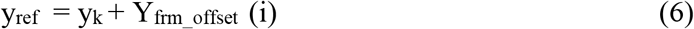

and its ROI reference strip can be determined by [x_roi_, y_roi_, w_roi_, h_roi_], where w_roi_ equals to the width of the synthetic reference frame, and

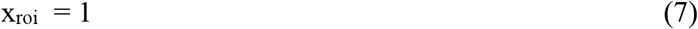

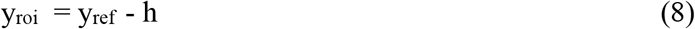

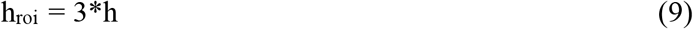

then we calculate the NCC using this strip and its corresponding ROI reference strip to identify the best matching location, and get its strip offset to represent the eye motion. In those rare occasions when two peaks were identified we used the one closest to the frame offset as the strip offset. We repeat this operation on all the frames to extract the strip offsets for the entire image sequence.

### Performance assessment

#### Comparison to benchmark

Since no ground-truth of the motion of the eye exists for our datasets, we chose a few different ways to evaluate its performance. One approach we took was to compare our new method to the registration method of Yang et al.^9^ that we considered for the purposes of this report to be our gold-standard benchmark. Unfortunately, due to fundamental differences between the two techniques, this comparison is imperfect and has several drawbacks (see discussion).

In our first attempted comparisons, we evaluated the results of each algorithm using comparable user-defined variables that govern the strip-wise registration, including strip size, number of strips and strip rejection threshold. As our studies of fixational eye movements required motion traces at a nominal temporal sampling rate of at least ∼480 Hz, we developed our approach using16 strips per frame and a strip height of 32 pixels and used these settings in each algorithm for our initial tests. However, we found that these parameters often caused the benchmark method to perform much more poorly than it did with its default parameter set. So, we tested a range of values and settled on using the default settings in the implementation of the offline digital registration version we had access to, that was implemented with a default of 15 strips per frame and a strip height of 64 pixels as this empirically gave the best results. The strip rejection threshold also differed as we used a variable threshold (see above), while the benchmark used a fixed threshold of 0.75.

To compare registration methods, we evaluated several aspects of the results, including: 1) the proportion of successfully registered data; 2) the standard deviation of pixel intensity in the registered image sequences; and 3) the energy of the high spatial frequency information in the final registered and averaged image. We also assessed structural repeatability in the registered and averaged images by comparing the variability in the spatial relationship between image features in the resulting images when different starting reference frames were manually selected.

The proportion of tracked frames or strips was computed by dividing the count of successfully tracked frames by the total number of frames in the sequence after excluding the blinks. It should also be noted that blinks are detected differently in the benchmark algorithm. For our eye tracking work using TSLO, we excluded from analysis discontinuities in motion traces of less than a single frame, so compare the proportion of tracked frames here. For AOSLO, we compare the proportion of tracked strips.

The variation in pixel intensity between the registered an unregistered image sequences was evaluated based on the hypothesis that after registration each pixel remains fixed on the same structure and thus experiences less variability in intensity over time than in the raw data where each pixel continuously sweeps across different structures. To assess this, we computed the standard deviation (SD) of each pixel across time; this was compared between the original and registered image sequences. This was done for cropped regions of each of the registered image sequences that excluded the margins of the image that may have had few strips contributing to each pixel. In addition, we also evaluated the SD of a registered image sequence that was composed only of frames that were dropped by the benchmark method but successfully tracked by the present approach. This allowed us to evaluate whether the additional frames tracked by the present approach were registered to the same level as those that were tracked by both.

The image energy of the high spatial frequencies in the image was computed using the registered and averaged images produced from each algorithm through the following steps: Each averaged image was filtered using a high pass filter with a normalized cutoff frequency of 0.02; the spatial variance was computed. This metric informs about the amount of energy in high spatial frequency features and therefore its value will decrease as image blur increases^22^. Finally, we computed the difference between the image energy generated from both registration methods. We plotted this difference divided by the energy of averages registered with our method in order to show the difference in energy as a percentage of the total energy of the averages.

#### Manual landmark-based performance evaluation

Finally, we also employed a manual method to evaluate whether the algorithm was appropriately registering image features that were easily identifiable by eye in the image sequence. To do this, we randomly picked 20 frames from each image sequence and manually marked the same features that were easy to identify in the images (e.g. vessel crossings). We selected several landmarks for each frame (see Fig.4). Multiple landmarks were chosen across the field of view to evaluate strip level accuracy across the entire image and to ensure that as the eye motion moved the field of view from frame to frame, that some landmarks would be visible in each image in the sequence. The manual marking was carried out by two independent graders, to allow us to compare the different human graders to one another and to the different registration algorithms. Bland-Altman^20^ plots were generated to evaluate the agreement between the different methods.

**Figure 4.**
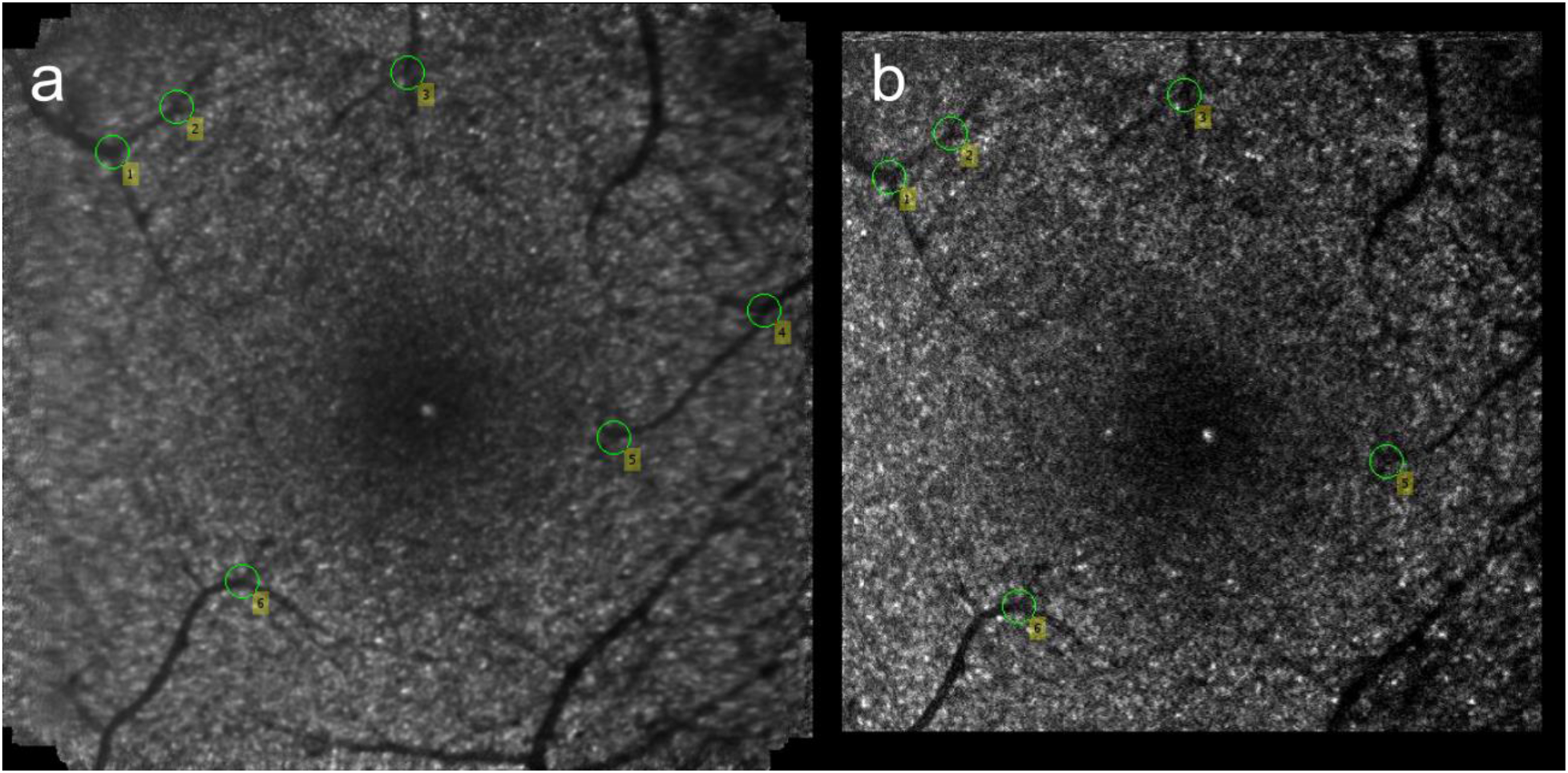
Manual landmarking. Several vessel landmarks were marked manually in Matlab. Screenshots from the marking script are shown here with several landmarks marked on the synthetic reference image (a) and then marked in an example target frames (b). This manual landmarking allowed for a direct comparison between the locations of readily identifiable image features marked by manual graders and their positions determined algorithmically.

#### Structural repeatability across different manually selected reference frames

To evaluate the structural repeatability of the final registered and averaged images, we compared the results of our algorithm to the benchmark when different reference frames are selected manually as the starting reference. For this, we used AOSLO image sequences and generated registered and averaged images for several starting reference frames. We then co-registered all of the averaged images generated from each algorithm, using sub-pixel registration^21^, and compared the spatial arrangement of the features across reference frame for each algorithm. These images were assessed both by visually evaluating animations generated from the registered images and by carefully comparing the positions of individual cone photoreceptors across the different images.

## RESULTS

### Comparison to benchmark

Across image quality levels in our TSLO datasets, our method successfully tracked 99.79% of all non-blink frames, leaving only 0.21% of all non-blink frames not successfully tracked. In comparison, the benchmark successfully tracked 68.89% of all non-blink frames across all quality levels, leaving 31.11% of frames untracked, on average. Table 1 lists the results for each of the different quality levels across the TSLO datasets and demonstrates that there were not major differences in performance across the range of image quality levels tested for the present algorithm with the proportion of unsuccessfully tracked frames ranged from 0.04–0.43%. In comparison, the benchmark was unable to track between 20.31 and 37.29% of the frames across each quality level.

**Table 1.**
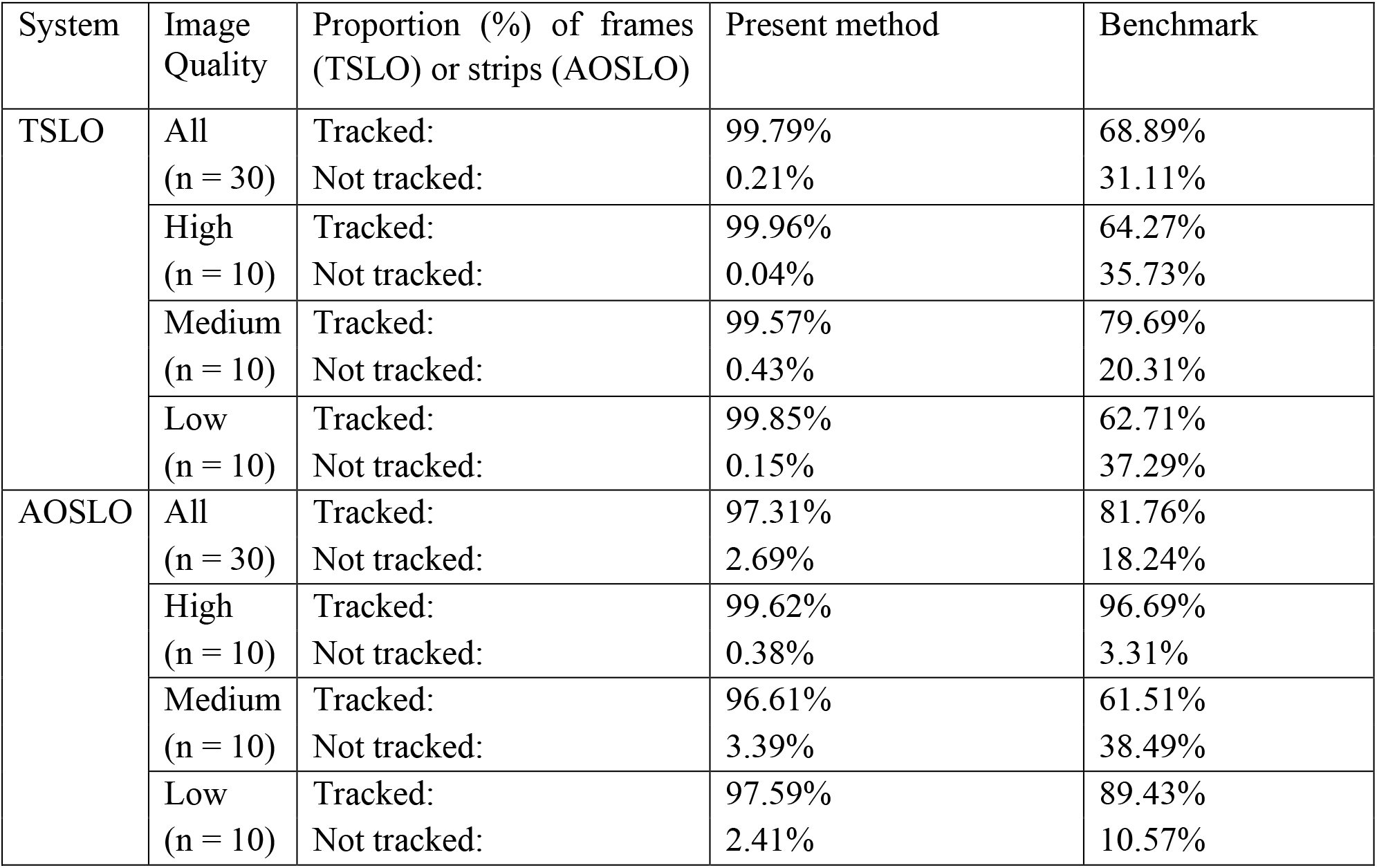
Proportion of tracked frames (TSLO) or strips (AOSLO) calculated excluding blinks. The proportion of tracked frames was similar across the range of image qualities tested for both the TSLO and AOSLO datasets. A larger proportion of frames were not successfully tracked by the benchmark algorithm across all test sets

Across image quality levels in our AOSLO datasets, our method successfully tracked 97.31% of all non-blink frame strips, leaving only 2.69% of all non-blink frame strips not successfully tracked. In comparison, the benchmark successfully tracked 81.76% of all non-blink frames across all quality levels, leaving 18.24% of non-blink frames untracked, on average. Table 1 lists the results for each of the quality levels across the AOSLO dataset. Again, there were not major differences in performance across the range of the image quality levels tested for the present algorithm; the proportion of unsuccessfully tracked frames ranged from 0.38–3.39%. In comparison, the benchmark was unable to track between 3.31 and 38.49% of the frames across each quality levels. Figure 5a shows the histogram of the SD across time for each pixel in the original data, 5b and c show histograms of the SD across time for each pixel in the registered image sequences created for the frames that were successfully registered by both algorithms; there is less variability in intensity over time in the registered image sequences than in the original raw data; in addition, there is very little difference in the distribution of each as is demonstrated by the difference in 5e. Similarly, 5d displays the histogram of SD across time for each pixel in the frames that were successfully tracked by our algorithm but that were not tracked by the benchmark; again, the distribution is very similar with the difference histogram in 5f showing very little difference between the two distributions.

**Figure 5.**
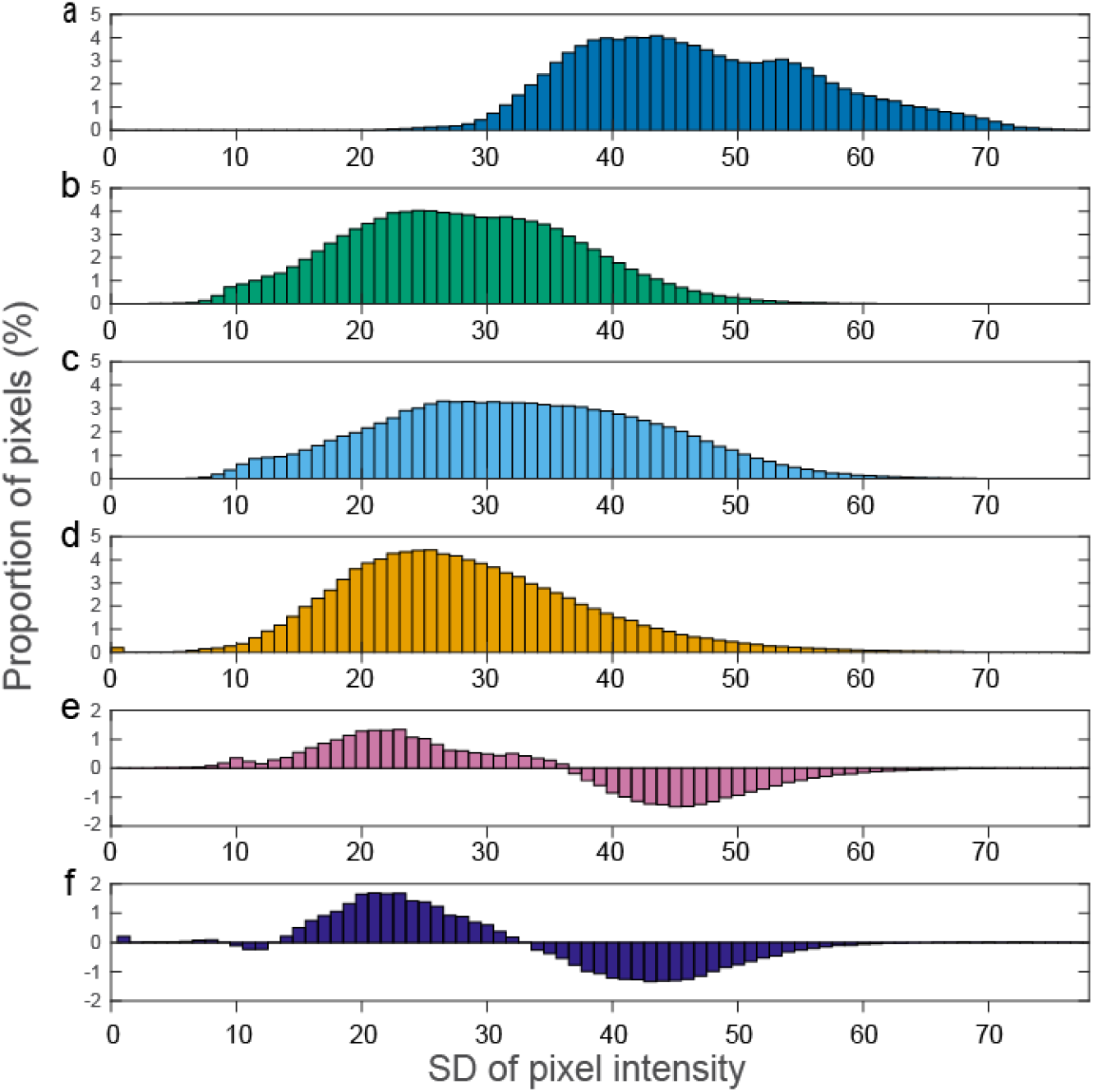
Standard deviation of pixels across time is reduced similarly for frames successfully tracked by each algorithm. The distribution of pixel SD across time for the (a) original raw data, for the registered data with the (b) current method and (c) the benchmark for frames successfully tracked by each, and (d) for the registered data only successfully tracked by our current method. (e) is the difference between distributions (b) and (c) showing that the distribution was similar for these two methods for frames successfully tracked by each. (f) is the difference between the distributions (c) and (d), showing that there was not a substantial difference in the proportion of pixels successfully tracked between the shared frames (e) or the frames only successfully tracked by the current method (f).

### Landmark comparison

We compared the manual landmarks made between the two graders and compared the differences between the graders and the algorithms; we also evaluated the agreement between the two algorithms. Bland-Altman plots evaluating the agreement between these different comparisons are provided in Supplementary Figure 2. The mean difference between the manual marking by different graders was less than a pixel (Suppl. Fig. 2a; −0.09 px horizontally and −0.27 px vertically). This average difference was smaller than the average difference obtained between the two algorithms that averaged 1.75 px horizontal and 1.5 px vertical (Suppl. Fig. 2b). Comparison between the manual graders and the two algorithms are shown in Suppl. Fig. 2c and Suppl Fig. 2d. The average difference between landmark method and our algorithm was a fraction of a pixel: −0.44 px horizontal and −0.23 px vertical, while there was a larger average difference for the benchmark of 2.193 px horizontal and −1.731 px vertical.

### Comparison of high spatial frequency information in registered and averaged images

We computed the high energy spectral band for each of the 30 registered and averaged images obtained in both TSLO and AOSLO. Figure 6 shows the normalized difference between these energy values across the images. For TSLO, there is a trend of increasing difference in energy between the two methods as image quality decreases. The inset shows two examples for two registered images that fall on different sizes of the quality spectrum. For the high quality image sequence there is no discernable difference in the resulting averaged image (pink arrow and pink outlined images) while the averaged images computed for the low-quality image sequence shows a much sharper and higher contrast image (green arrow and outlined images). This qualitative result is borne out in the energy metric. For the first example the difference in energy is almost none while for the second it is almost 75%.

**Figure 6:**
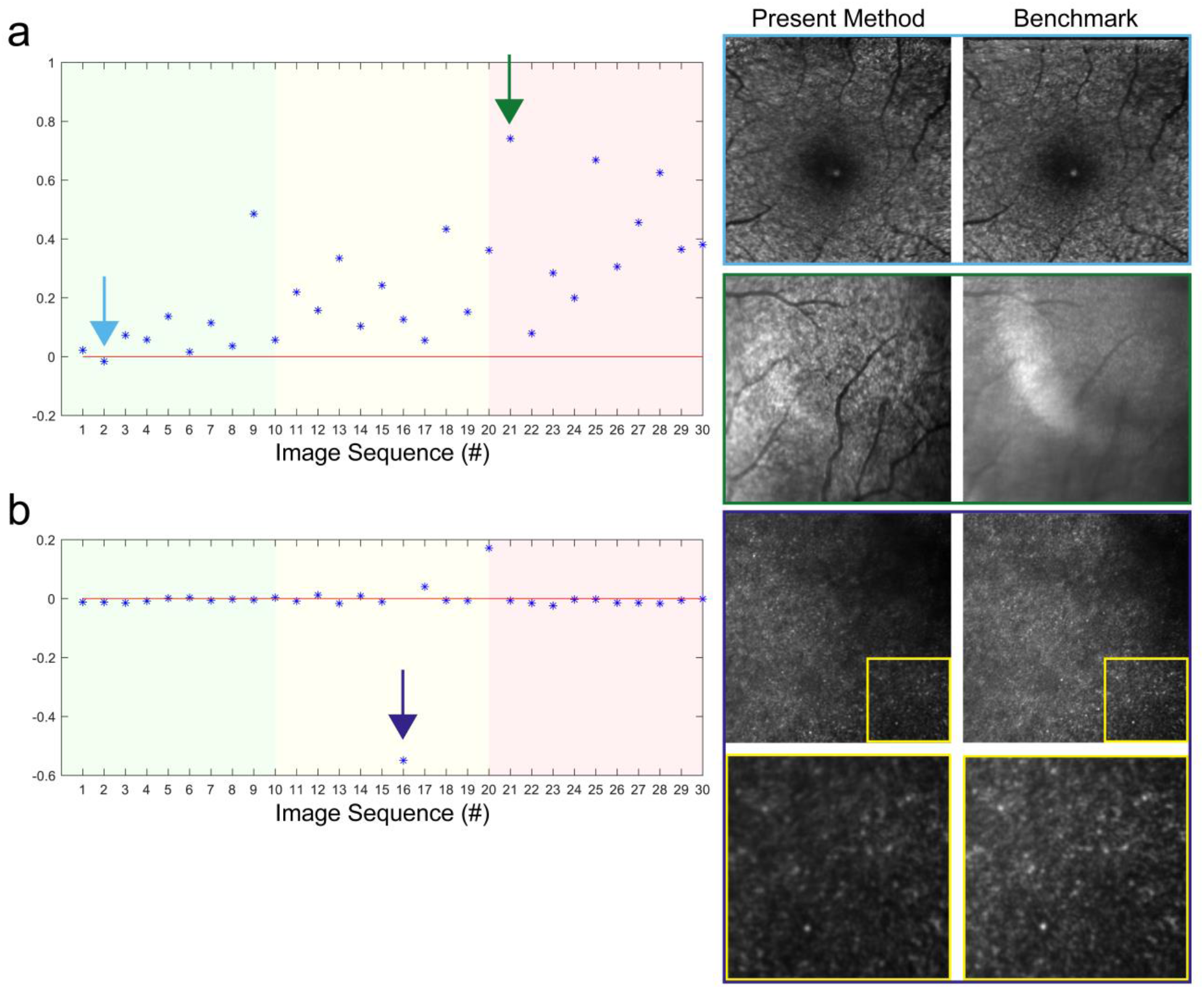
Normalized difference in energy of high spatial frequencies between our method and the benchmark for the TSLO (a) and AOSLO (b) image sequences. Background color denotes subjective quality of the image sequence (high = green; medium = yellow; red = low). TSLO datasets showed a range of differences, with small differences resulting in negligible differences in subjective image quality (e.g. light blue arrow in (a) and corresponding images traced in same color to the right). There were larger differences for lower quality image sequences and the present method produced images with subjectively higher quality than the benchmark (e.g. dark green arrow in (a) for corresponding averaged images traced in same color to the right). The difference was very small for most AOSLO image sequences aside from one outlier that demonstrated better subjective image quality from the benchmark compared to the present method (purple arrow in (b) and images to right traced in corresponding color). This is appreciated better by examining the zoomed in section (bottom images traced in yellow, location denoted by yellow squares in the larger images above).

The same metric applied to the AOSLO image sequences is shown in Fig.6 (b). For this layer (i.e. photoreceptor layer) with this instrument both algorithms have a similar performance as the difference in energy is mostly close to zero except for two exceptions. For image sequence number 16 our algorithm seems to underperform with respect to the benchmark. A region of the averages on the right side of the plot is enlarged showing how the photoreceptors from the image produced by our method are indeed blurrier than the ones on the image obtained with the benchmark.

### Structural repeatability

For AOSLO, we also evaluated the structural repeatability of the images by comparing the resulting averaged images when the registration was seeded with different manually selected reference frames. The differences in the spatial position of image features is best appreciated by evaluating the animation shown in multimedia item 1. This animation shows the averaged images, co-registered for each method, side-by-side for a representative high quality AOSLO image sequence when 5 different reference frames were manually selected. It should be noted that each reference frame appeared to be free from obvious eye motion distortions and considered to be of equivalent subjective image quality. This animation shows that there is little variation in the spatial arrangement of image features for our method, while the images obtained from the benchmark method show variations in the positions of each cell from image to image. This variation in cell position is evaluated for several cells across these images in Figure 7.

**Figure 7.**
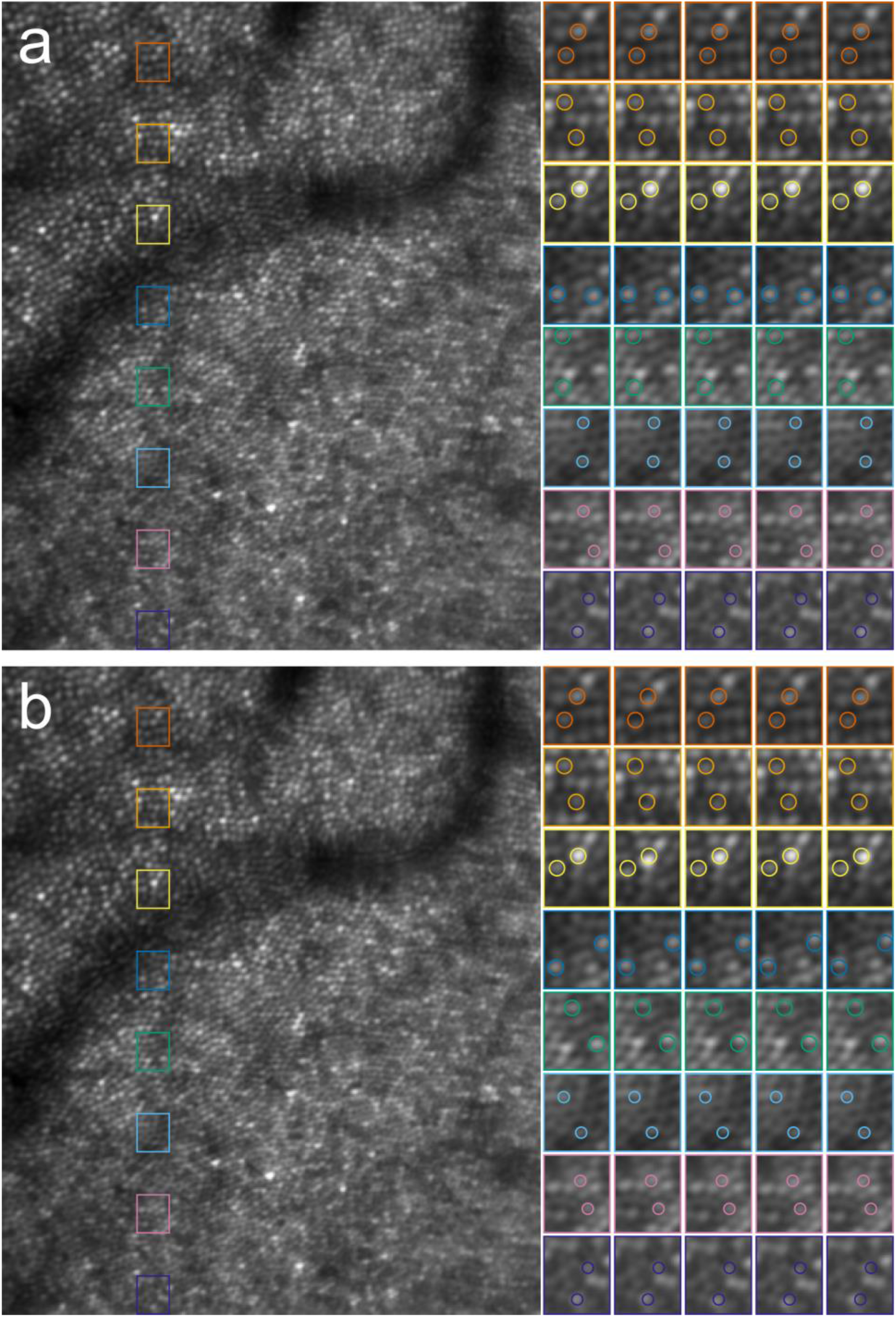
The same spatial arrangement of cells is seen no matter what reference frame is selected. The present method (a) and benchmark (b) were used to generate averaged images when starting with the same five different reference frames. The full averaged image generated when using the first reference frame is shown at left. The overlaid colored rectangles in each denote the positions of the location of the zoomed in locations shown to the right for each of the different reference frames. Two cones were circled in the first image (zoomed images, left column) and then overlaid at the same location on the other images to evaluate cell location repeatability. Nearly all cones for the present method remained in the same location no matter what reference was used. This was not the case for the benchmark, where many cones (e.g. orange, yellow, light blue rectangles) shifted positions depending on the manual reference frame selected. Differences are small but can be on the order of a whole cone and are most apparent when toggling between images (see animation).

Here we show the resulting image for the first reference frame shown in the animation (for reference frame 1) on the left side. The 8 colored squares arrayed vertically across the image denote the location of the zoomed in views of those regions that are shown for each of the five reference frames in the colored rectangles to the right. Within each of these small regions of interest that are shown with 2x magnification to the right we have denoted the position of two of the cones in the first image with colored circles. The location of these cones in the image generated using the first reference frame is overlaid on the other four images to show whether that cone remained in the same location across references or whether it shifted position. Nearly every cone was in the same location for each of the cones evaluated in the images generated from our method. There are some small shifts that can be appreciated in the animation and in a few instances here we see some small variability in cone position across reference frames (e.g. slight shifts for the lower cones in the yellow region and in the light blue region for images 3-5). The variability in cone position was greater for the benchmark method and could result in position differences across the different images on the order of the size of an individual cone (e.g. orange, green, and cyan areas).

## III. DISCUSSION

We have shown here that when time constraints are no longer a limiting factor, additional image processing steps can be applied to achieve an improved registration result, both for eye tracking and for imaging applications. Key aspects of our approach include a pre-processing contrast enhancement phase and routines for detecting distorted and candidate large motion frames. Perhaps one of the most different aspects of our approach compared to what has been published previously in this area is the development of a composite reference frame for the final strip level registration, however, it should be noted that this approach was proposed previously by Stevenson. This work builds on the previous achievements of Stevenson, Roorda, Yang and others to demonstrate that certain aspects can be improved when the time constraints imposed by real time tracking are lifted.

Registration algorithms designed for eye tracking are inherently difficult to assess without a ground truth reference for comparison. This limitation forced us to utilize approaches for assessment that had limitations. For one, there were inherent differences between our method and the comparison algorithm we chose as our gold-standard benchmark that do not always put them on equal footing. Differences of note here are the different number of strips used, the different blink detection method used and the different size of the strips, etc. It was our original intention to set all parameters to be identical, however, when doing so we found that this would not reflect the performance that could be achieved with the benchmark algorithm when using its default settings, so we chose to use those parameters instead.

In terms of proportion of tracked frames or strips, we showed that our technique could achieve very high tracking rates, approaching 99% in most cases. This reflected a substantially greater proportion of data that could be successfully tracked compared to the benchmark, so we were interested to know if those additional strips were tracked with the same level of precision as those that could be successfully tracked by both techniques. Comparison of the registered image sequences between algorithms using the SD across time method showed that when frames were successfully registered in each algorithm, a similar reduction in SD across time was seen in the registered image sequence (Fig.5). This demonstrates that the additional frames tracked by our method are registered to a similar level of accuracy at least as it is reflected by this metric. Careful inspection of the difference histograms shows that there was a small shift in the distribution towards slightly higher SD in the additional frames successfully tracked by our algorithm that were dropped by the benchmark (Fig. 5e). This small difference could reflect a decrease in the computation accuracy or that these frames display a higher variation in overall intensity due to large eye motion.

Improved strip tracking is beneficial for eye tracking applications, but it should be noted that keeping more frames does not always improve the quality of the final registered and averaged image for imaging applications. In certain cases, averaging more strips/frames can have a detrimental effect on image quality. The need to include more data in the registered and averaged image is most important for light starved imaging applications like autofluorescence and some forms of non-confocal AOSLO. For example, confocal AOSLO imaging of photoreceptors may only require a relatively small number of strips per pixel to be averaged to achieve a high-quality image. In those cases, one likely would only include in the final averaged images those strips that had the highest cross-correlation threshold. For those applications we have generated an averaging and cropping tool that allows the registered image sequences averaging to be customized to the application.

However it should be noted that in the cases where the registration algorithm fail to provide accurate offsets measures, it leads to blurry average images as shown in Fig.6. We show how the degradation of the video quality is linked to a lower value of the energy of high frequency structures and to a blurrier image for the benchmark method compared to ours. The video quality can decrease because patient cannot fixate properly due to their disease or even fatigue which can often occur in a clinical environment and is therefore important for registration algorithm to achieve good accuracy in these instances.

In addition, a poor estimation of frame motion like the one shown in Fig.6 for some of these TSLO videos prevented us from undertaking studies requiring the accurate tracking of the eye fixational movements such as our work evaluating their correlation with concussion^16^.

For evaluating the tracking precision, we developed a tedious manual landmarking approach and deployed it on several image sequences. This was a suitable method for the TSLO data where larger vessel landmarks could be reliable marked by our human graders, but it was not useful for the AOSLO data, as we found the additional structural image detail in AOSLO made it nearly impossible for the manual graders to reliably mark the same exact structure from frame to frame. Despite this limitation, we showed that human graders could routinely detect the same structures reliably and that there were larger differences between the different algorithms than there were between the different graders.

The ability to track single cells across time within the living eye has long been an overarching goal for AO Ophthalmoscopes. However, we have been limited in our ability to track cells longitudinally due to our inability to reliably reproduce the same retinal structure in the face of within frame distortions from eye movements. Some investigators have taken the approach of warping the averaged images that they want to compare across images within an imaging session or across images taken at different timepoints. This is suitable for psychophysical testing or imaging studies on normal eyes when the retina is not expected to change between imaging sessions. However, this is unsuitable when the retinal structure is changing such as in progressing disease or in response to treatment. The ability we have shown here to reliably reproduce the same retinal structure (Fig. 7) will facilitate all imaging applications as in all cases we seek to recapitulate the true arrangement between structures within the imaging field of view. However, it is the evaluation of cell and gene-based treatments that we think will benefit the most from this precise level of targeted cell tracking.

Despite our achievements there are several aspects that could still be improved further. For one, our blink detection approach is simplistic and fails when there are not blinks in an image sequence. This is an area in certain need of improvement. Other aspects that could be improved would be to implement an automatic reference frame selection step. We plan to do this to make the pipeline more automated; this can be done using any of several image quality metrics, as others have done previously. Finally, the main drawback in our approach is that it takes a long time in its present implementation to process the data. At present, using CUDA implementations in Matlab only, it takes approximately 9 minutes to process 900 frames of data using the parameters outlined above. However, this long computational time can be reduced through software modifications, such as porting computationally intensive tasks to other languages or using additional hardware.

## CONCLUSIONS

We show here that our modifications to the strip-based digital image registration approach for scanned ophthalmic imaging systems accomplished our primary objectives:

1. Tracks the precise motion of nearly all the images in each sequence for eye-tracking and light starved imaging applications.
2. Is sensitive to motion larger than the field of view of a single frame.
3. Reconstructs the spatial arrangement between images consistently and accurately.

Taken together, these improvements extend the current capabilities of strip-based digital image registration for eye tracking and imaging applications. Our technique facilitates the study of fixational eye movements, an emerging area of importance for understanding early changes in diseases of the eye and brain. It will also enable the tracking of individual cells over time in health and disease to permit targeted monitoring of individual cells in response to treatment.

