## Supplemental Figure 1 for "Improvements to strip-based digital image registration for robust eye-tracking and to minimize distortions in images from scanned ophthalmic imaging systems"

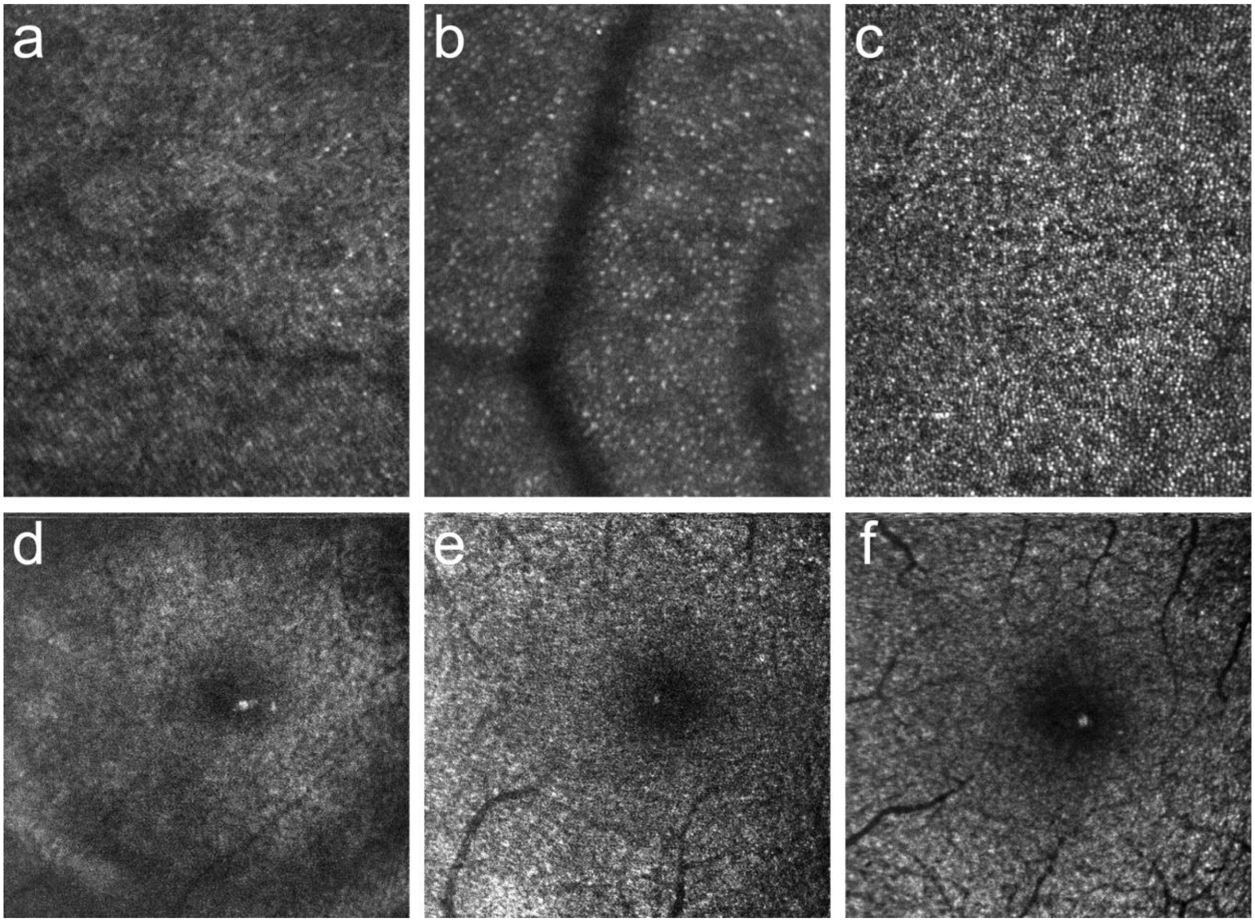


**Supplemental Figure 1. Representative examples of range of image quality in test data.** Top row shows examples from AOSLO of low (a), medium (b) and high quality (c) frames, while bottom row shows example of low (d), medium (e) and high quality (f) frames for TSLO data. Image height is 1.5° for AOSLO images and 5° degrees for TSLO.
