## Supplemental Figure 2 for "Improvements to strip-based digital image registration for robust eye-tracking and to minimize distortions in images from scanned ophthalmic imaging systems"

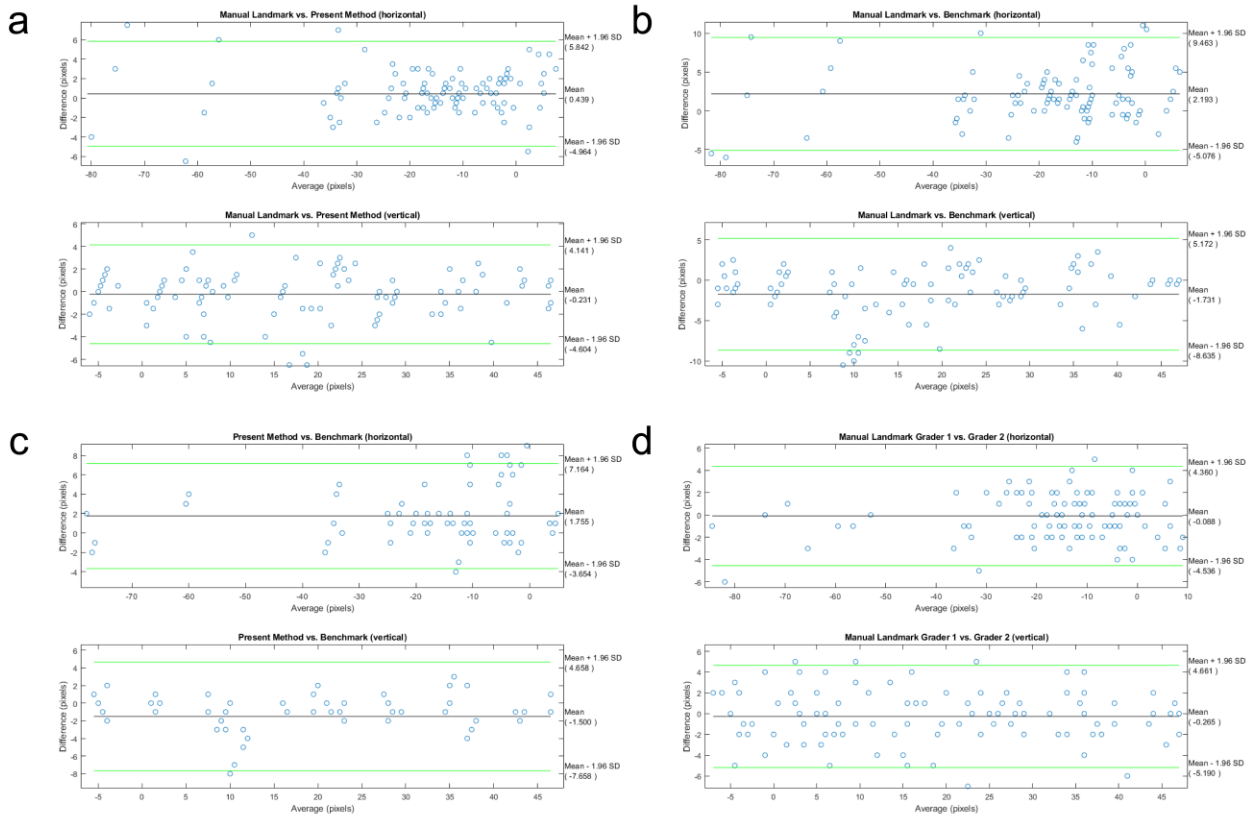


**Supplemental Figure 2. Bland-Altman plots show agreement between present method, benchmark and manual graders.** The average difference was less than 0.5 pixels in either direction between the manual landmark and present method (a), while the difference was larger, around 2 pixels, on average, for the manual landmark versus the benchmark (b). There was also little difference seen when comparing the positions between the present method and the benchmark (c). The agreement between the two graders was better than any other comparison (less than 0.3 pixels difference, on average).
